# Relative Homoplasy Index: A New Cross-comparable Metric for Quantifying Homoplasy in Discrete Character Datasets

**DOI:** 10.1101/2023.10.10.561677

**Authors:** Elizabeth M. Steell, Allison Y. Hsiang, Daniel J. Field

## Abstract

Homoplasy is among the main hinderances to phylogenetic inference. However, investigating patterns of homoplasy can also improve our understanding of macroevolution, for instance by revealing evolutionary constraints on morphology, or highlighting convergent form-function relationships. Several methods have been proposed to quantify the extent of homoplasy in discrete character matrices, but the consistency index (CI) and retention index (RI) have remained the most widely used for decades, with little recent scrutiny of their function. Here, we test the performance of CI and RI using simulated and empirical datasets and investigate patterns of homoplasy with different matrix scenarios. In addition, we describe and test a new scaled metric, the relative homoplasy index (RHI), implemented in the R statistical environment. The results suggest that, unlike the RI, the CI does not constitute a direct measure of homoplasy. However, the RI consistently underestimates the extent of homoplasy in phylogenetic character-taxon matrices, particularly in datasets characterised by high levels of homoplasy. By contrast, RHI—the newly proposed metric—outperforms both methods in sensitivity to homoplasy levels, and is scaled between zero and one for comparison of values between different datasets. Using both simulated and empirical phylogenetic datasets, we show that relative levels of homoplasy remain constant with the addition of novel characters, and, in contrast to earlier work, decrease with the addition of taxa. Our results help illuminate the inherent properties of homoplasy in cladistic matrices, opening new potential avenues of research for investigating patterns of homoplasy in macroevolutionary studies.

## Introduction

Homoplasy is an inevitable property of discrete character matrices constructed for phylogenetic inference. In cladistic matrices, homoplasy is defined as the independent acquisition of or reversal to the same character state (Lankester 1870; Sanderson and Donoghue 1989), and is often considered a limitation in morphological datasets as it can obscure phylogenetic relationships (Archie 1996). However, with ever-improving methods to infer phylogenetic relationships and the increasing availability of genomic data for extant clades, accurately quantifying the extent of homoplasy in phylogenetic datasets has potential to be useful for informing estimates of evolvability and constraints on morphological evolution (Sanderson and Donoghue 1989; Wagner 2000; Sidlauskas 2008; Brocklehurst and Benson 2021).

Methods to estimate homoplasy received considerable attention in the late 20^th^ century with an extensive body of research exploring parsimony-based metrics with cladistic matrices (summarised in Archie 1996). In recent years, however, few studies have explored patterns of homoplasy in discrete datasets, and the development of new homoplasy estimation methods have lagged behind methodological advances in phylogenetic inference. The two most widely-implemented and easily calculated homoplasy metrics are the parsimony-based consistency index (CI) (Kluge and Farris 1969) and retention index (RI) (Farris 1989). These metrics are implemented in phylogenetic inference programmes such as TNT (Goloboff et al. 2008) and the ‘phangorn’ *R* package (Schliep 2011), and have been subject to numerous comparative analyses on their efficacy for detecting homoplasy, and used to explore patterns of homoplasy within discrete character matrices (Sanderson and Donoghue 1989; Meier et al. 1991; Archie 1996; Hauser and Boyajian 1997; Sookias 2020; Murphy et al. 2021).

However, CI and RI are not cross-comparable methods; that is, they cannot be directly compared across alternative datasets and phylogenetic trees, limiting attempts to infer generalisable conclusions about patterns of homoplasy in phylogenetic datasets. For instance, CI is highly sensitive to variable numbers of taxa (but not characters) in phylogenetic matrices, with a negative relationship detected between CI and the number of taxa included (Sanderson and Donoghue 1989; Hauser and Boyajian 1997; Murphy et al. 2021). By contrast, RI represents a more robust metric with respect to matrix dimensions (Hauser and Boyajian 1997; Murphy et al. 2021). Nonetheless, values of RI are not scaled between zero and one; instead, the minimum value (which indicates the maximum amount of homoplasy possible) is an arbitrary value somewhere in that range (Archie 1996). This property arises because RI is calculated using the theoretical maximum treelength for a given dataset; that is, the number of steps estimated for a matrix when the tree is a complete polytomy (Farris 1989; Archie 1996). Therefore, in any dataset where the tree has more than one resolved node, RI will never be truly scaled between zero and one, rendering values of RI non-comparable between different datasets. To overcome this problematic issue, Archie (1989) developed the homoplasy excess ratio (HER). Unfortunately, however, this method has its own limitations in that it is not applicable to phylogenies inferred under optimality criteria other than maximum parsimony. For instance, phylogenies inferred through Bayesian inference or incorporating topological constraints are not suitable for this method, as HER implements maximum parsimony to infer topologies from randomised versions of character matrices in order to estimate an expected maximum value of homoplasy. Furthermore, HER is particularly time consuming and computationally demanding to calculate, and as such has not seen wide adoption.

Towards improving the rigour of systematic investigations of patterns of homoplasy, we present a novel metric, the relative homoplasy index (RHI), for estimating the extent of homoplasy in phylogenetic datasets. The algorithmic basis for RHI is similar to RI, but the resulting metric is consistently scaled between zero and one for empirical datasets, thus facilitating statistically meaningful comparisons across datasets with differing numbers of taxa and characters. We compare RHI to CI and RI using simulated matrices and empirical datasets, and test the sensitivity of each metric to variation in numbers of taxa, characters, and character state transition rates. We also investigate six empirical datasets deriving from four recent studies on bird phylogenetic relationships (Ksepka et al. 2019; Field et al. 2020; Benito et al. 2022; Steell et al. 2023) to explore these patterns of homoplasy in various bird clades, and analyse simulated character matrices based on these empirical topologies to account for systematic bias from random tree generation methods.

## Materials & Methods

### Homoplasy Indices

*Consistency index—*The consistency index (CI) is defined as:

(1)

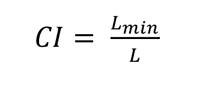

where *L* represents tree length (the number of steps for a tree with a given matrix, regardless of whether the tree was inferred from that matrix or not) and *L_min_* represents the theoretical minimum tree length for a matrix with the same dimensions (i.e., same number of taxa, characters, and proportion of multistate characters) – that is, the length of the tree if there were no homoplasy in the dataset (Kluge and Farris 1969; Farris 1989). *L_min_* is calculated by:

(2)

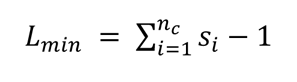

where *s_i_* is the number of states for the *i^th^* character, and *nc* is the total number of characters in the matrix. As *L_min_* is necessarily always ≤ *L* by definition, CI is always a value between 0 and 1 (0 < CI ≤ 1; *L_min_* cannot equal 0 as long as there is at least one non-invariable character in the matrix). CI is generally interpreted as how well a dataset ‘fits’ a particular tree under the assumption of parsimony (the ‘consistency’; Kluge and Farris 1969). The complement 1 – CI can then be interpreted as how poorly a dataset fits the tree under consideration.

*Retention index*—The retention index (RI) is defined as:

(3)

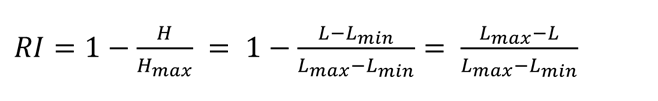

where *H* represents the total number of extra steps (i.e., steps due to homoplasy; Kluge and Farris 1969; Farris 1989), and can be calculated by subtracting *L_min_* from the observed tree length *L*; *L_max_* represents the theoretical maximum tree length for the matrix, which is the tree length when the matrix is applied to a completely unresolved tree (star tree or single polytomy) (Farris 1989; Archie 1996; Cuthill et al. 2010).

CI and RI can be expressed with respect to each other:

(4)

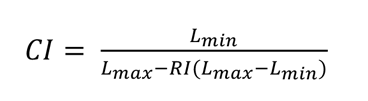

(5)

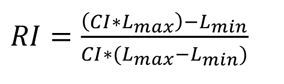

From these equations, it follows that RI is equal to 0 when CI = L_min_/L_max_, i.e., when L = L_max_.

*Relative homoplasy index*—We introduce a novel metric, the relative homoplasy index (RHI), which is defined as follows:

(6)

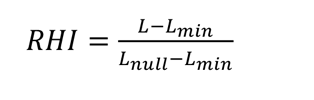

where *L_null_* (null tree length) represents the tree length for the matrix when no phylogenetic signal is present except that which is present due to chance. *L_null_* is calculated by taking the median tree length for the empirical matrix with a set of randomised trees. In these randomised trees (null trees), the topology of the empirical tree (i.e., tree shape) is maintained, but the tip names are randomised. Therefore, the properties of the empirical tree—such as number of supported nodes, tree balance, and tree shape—are retained in the set of null trees, avoiding systematic biases associated with these properties (Felsenstein 2004; Simmons et al. 2004).

The relative homoplasy index differs from the consistency index and retention index in that 0 indicates an absence of homoplasy for a matrix (when *L* = *L_min_*), and 1 indicates that a matrix shows as much homoplasy as possible when the corresponding topology is resolved beyond a polytomy (when *L* = *L_null_*). In this case, a value of 1 would occur if the matrix contains no phylogenetic information with respect to the tree, beyond that which is present by chance. RHI can be calculated for any discrete matrix with a given phylogeny, regardless of the method or dataset that the phylogeny was inferred from. RHI is a cross-comparable metric that can be applied to matrices with accompanying phylogenetic trees that may have been generated from independent matrices (e.g., molecular data), when the topology is a consensus between multiple datasets, or when the topology is partially constrained by another phylogenetic hypothesis.

### Data Acquisition and Analyses

We selected published phylogenetic datasets with a focus on birds to investigate homoplasy in this study. Datasets were selected based on their properties such as tree inference method, taxon and character dimensions, and availability of data from public repositories. The six datasets used in this study are summarised in Table 1 and were taken from four published studies with morphological matrices of different bird taxa. Datasets from Benito et al. (2022) and Steell et al. (2023) were formatted so that two different treatments of the matrix and/or empirical tree could be analysed. For Benito et al., two datasets were analysed (Avialae *a* and Avialae *b*). Both matrices are the same, but the topologies are based on different phylogenetic inference methods; the topology for Avialae *a* was inferred under maximum parsimony, and the topology for Avialae *b* was inferred under Bayesian inference. For Steell et al., two datasets were analysed with the same characters but slightly different taxon samples and topologies (Passeriformes *a* and Passeriformes *b*). Passeriformes *a* includes only extant taxa, and the topology was not inferred from the morphological matrix, but from entirely independent molecular datasets from other studies (Prum et al. 2015; Oliveros et al. 2019; Harvey et al. 2020). Passeriformes *b* includes all taxa from Passeriformes *a* in addition to the fossil specimens analysed in the original publication, with the topology inferred from the morphological dataset with a topological constraint for the extant taxa based on the aforementioned molecular studies. Empirical topologies used in this study are represented in Figure 1. The topologies used in this study were taken as previously published; as such, some are unrooted and all except Passeriformes *a* are incompletely resolved (Fig. 1; Table 1). We chose to leave the trees in this condition instead of reanalysing them as we were interested in investigating patterns of homoplasy across a range of real datasets and phylogenetic trees, as opposed to reanalysing phylogenetic relationships.

**Figure 1.**
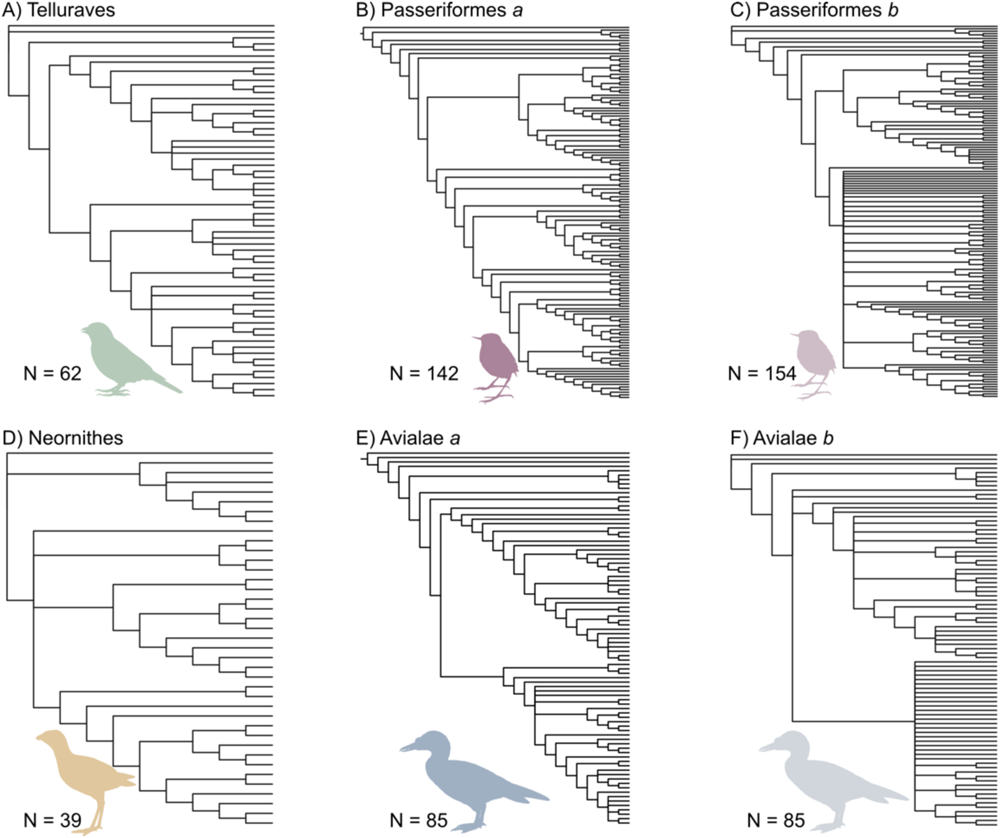
Phylogenetic trees without branch lengths used in analyses. A) Unrooted, partially constrained topology of Telluraves from Ksepka et al. (2019); B) Rooted, completely constrained topology of Passeriformes a (extant taxa only) from Steell et al. (2023); C) Unrooted, partially constrained topology of Passeriformes b (extant and fossil taxa) from Steell et al. (2023); D) Unrooted, partially constrained topology of Neornithes from Field et al. (2020); E) Rooted, unconstrained Maximum Parsimony derived topology of Avialae a from Benito et al. (2022); F) Unrooted, unconstrained Bayesian derived topology of Avialae b from Benito et al. (2022).

**Table 1.** Summary of datasets.

| Dataset | Tree | Taxa | Chars. | Taxa:<br>chars. | State #<br>proportions<br>(binary: 3: >3) | Supported<br>nodes (%) | Missing<br>data<br>(%) |
| --- | --- | --- | --- | --- | --- | --- | --- |
| Telluraves <sup>a</sup> | MP | 62 | 156 | 0.40 | 0.81:0.23:0.03 | 82.0 | 27.5 |
| Passeriformes<br>(a) <sup>b</sup> | - | 142 | 49 | 2.90 | 0.86:0.14:0 | 100.0 | 7.5 |
| Passeriformes<br>(b) <sup>b</sup> | Bayesian | 154 | 49 | 3.14 | 0.86:0.14:0 | 80.4 | 11.1 |
| Neornithes <sup>c</sup> | Bayesian | 39 | 295 | 0.13 | 0.67:0.28:0.04 | 89.5 | 18.4 |
| Avialae (a) <sup>*</sup><br><sup>d</sup> | MP | 85 | 282 | 0.30 | 0.70:0.23:0.07 | 82.1 | 51.1 |
| Avialae (b) <sup>*</sup><br><sup>d</sup> | Bayesian | 85 | 282 | 0.30 | 0.70:0.23:0.07 | 45.2 | 51.1 |

\*No topological constraints applied in original phylogenetic analysis. Percentage of supported nodes is based on percentage of fully resolved nodes in the original publication of the phylogeny. <sup>a</sup> Ksepka et al. (2019); <sup>b</sup> Steell et al. (2023); <sup>c</sup> Field et al. 2020; <sup>d</sup> Benito et al. 2022. Character numbers may differ from the original publication because invariant characters were removed prior to analyses in this work.

All analyses were carried out in *R* v4.2.2 (R Core Team 2021), primarily using functions within the ‘APE’ (Paradis et al. 2004), ‘phangorn’ (Schliep 2011), and ‘dispRity’ (Guillerme 2018) packages. All code used here, including novel functions, can be found at github.com/LizzySteell/Homoplasy. Invariant characters were removed prior to analysis. Invariant characters are uninformative with respect to homoplasy, and removal was necessary because the RHI function requires the minimum number of states for any character to be two, rather than one. Inapplicable characters (usually represented as ‘-’) were treated as ambiguities (missing states; ‘?’) for simplicity. Using the ‘phyDat’ matrix format implemented in ‘phangorn’, we used contrast matrices to specify how steps should be counted for ambiguities and polymorphic scorings. Contrast matrices enable ambiguities (‘?’) to be interpreted as all possible numbered states (i.e., ‘0’, ‘1’, ‘2’, etc.), meaning no step is counted from a numbered state to an ambiguity and vice versa. Polymorphisms are interpreted as all numbered states specified in the polymorphism (i.e., ‘{01}’ will be interpreted as both ‘0’ and ‘1’). This means that when polymorphisms are present, the least costly state in the polymorphism contributes to the tree length (for further discussion, see Watanabe 2016). In this way, matrices in the ‘phyDat’ format implemented by ‘phangorn’ can be analysed with the same treatment of polymorphisms as in maximum parsimony packages such as TNT (Goloboff et al. 2008). Therefore, all metrics calculated here represent a *minimum* estimate of homoplasy, and increasing the proportion of ambiguities in a matrix will further decrease the amount of homoplasy detected.

Consistency index and retention index were calculated for all empirical datasets with all characters treated as unordered. CI and RI were calculated using CI() and RI() functions from ‘phangorn’. For illustrative purposes CI and RI values are reported in the figures as 1 – CI and 1 – RI, respectively, such that a value closer to 1 indicates more homoplasy than a value closer to 0. This allows for a scale that is consistent across metrics and more easily comparable. A new function was written to calculate the relative homoplasy index (RHI()) that includes the option to specify certain characters as ordered. The arguments for RHI() are ‘data’ (matrix read into R using read.nexus()), ‘tree’ (phylogenetic tree of class ‘phylo’), ‘n’ (number of null trees to evaluate), ‘contrast’ (object containing a contrast matrix), ‘ord’ (a vector specifying ordered characters; default = NULL) and ‘cost’ (object containing a cost matrix to specify the number of steps taken during transitions of ordered character states; default = NULL). To obtain a distribution of null tree lengths for RHI, tips were randomised using sample() without replacement while maintaining the overall topology (tree shape). The parsimony() function was then applied to the empirical matrix and each randomised tree to obtain a distribution of values for *L_null_*. Null tree length values are normally distributed (Supplementary Figure 1, Appendix 1) but the range of values varies depending on the properties of the supplied tree and matrix. Generally, we recommend *n* = 1000; but *n* = 100 may be adequate based on the simulations and the close similarity of the *L_null_* distributions at different values of *n* (Supp. Fig. 1). The function outputs the median *L_null_*, 5% and 95% quantiles, tree length (*L*) and theoretical minimum tree length (*L_min_*). The second output in the list returned by RHI() is a vector containing the null tree lengths.

We carried out three analyses using simulated matrices based on empirical topologies, and three analyses with empirical topologies and associated empirical datasets. Analyses are summarised in Table 2.

**Table 2.** Summary of analyses.

| Analysis | Datasets | Independent variable (X) | X values | Trees per run | Matrices per tree | Runs (# of X values) | Total iterations per run |
| --- | --- | --- | --- | --- | --- | --- | --- |
| *Simulated matrices* |  |  |  |  |  |  |  |
| 1. Inflating transition rate | Telluraves<br>Passeriformes (a)<br>Neornithes<br>Avialae (a) | Mean of log transformed transition rate ($\mu$) | -8 to 2 by 0.5 | 1 | 20 | 22 | 420 |
| 2. Varying number of taxa: sub-tree sampling | Telluraves<br>Passeriformes (a) | Number of taxa | 10 to 62 by 2<br>10 to 140 by 5 | 100 | 1 | 27<br>27 | 2700<br>2700 |
| 3. Varying number of characters | Telluraves<br>Passeriformes (a) | Number of characters | 10 to 145 by 5<br>10 to 48 by 2 | 1 | 100 | 28<br>20 | 2800<br>2000 |
| *Empirical matrices* |  |  |  |  |  |  |  |
| 4. Comparison of | Telluraves<br>Passeriformes (a) | na | na | na | na | 1* | 1000* |
| homoplasy metrics across datasets | Passeriformes (b)<br>Neornithes<br>Avialae (a)<br>Avialae (b) |  |  |  |  |  |  |
| 5. Varying number of taxa: sub-tree sampling | Telluraves | Number of taxa | 10 to 60 by 5 | 100 | 1 | 11 | 1100 |
|  | Passeriformes (a) |  | 10 to 140 by 10 |  |  | 14 | 1400 |
|  | Passeriformes (b) |  | 10 to 150 by 10 |  |  | 15 | 1500 |
|  | Neornithes |  | 10 to 38 by 2 |  |  | 15 | 1500 |
|  | Avialae (a) |  | 10 to 80 by 5 |  |  | 15 | 1500 |
|  | Avialae (b) |  | 10 to 80 by 5 |  |  | 15 | 1500 |
| 6. Varying number of characters: matrix sampling | Telluraves | Number of characters | 10 to 140 by 10 | 1 | 100 | 14 | 1400 |
|  | Passeriformes (a) |  | 10 to 48 by 2 |  |  | 20 | 2000 |
|  | Passeriformes (b) |  | 10 to 48 by 2 |  |  | 20 | 2000 |
|  | Neornithes |  | 10 to 290 by 20 |  |  | 15 | 1500 |
|  | Avialae (a) |  | 10 to 280 by 20 |  |  | 14 | 1400 |
|  | Avialae (b) |  | 10 to 280 by 20 |  |  | 14 | 1400 |

\* 1 run per metric and 1,000 iterations (n = 1,000) for RHI only.

### Investigating Homoplasy in Simulated Datasets

Matrices were simulated with topologies following those in the source publications (Fig. 1) for the six datasets (Table 1). These empirical topologies were selected as reference trees for simulations with the aim of simulating realistic character matrices. Random tree generation methods, such as the coalescent, birth-death or non-evolutionary model methods implemented in ‘APE’ (Paradis et al. 2004) each produce trees with characteristic properties, such as particular tree shapes, with different frequencies. Therefore, publication-derived empirical topologies were used to avoid potential systematic bias resulting from random tree generation methods. Estimating the extent of how homoplasy varies with respect to differing tree shapes is beyond the scope of this study.

Matrix simulation required fully resolved (bifurcating) trees with branch lengths. We used the accelerated transformation (ACCTRAN) algorithm (Farris 1970; Swofford and Maddison 1987) as implemented in phangorn’s acctran() function with each empirical topology and its corresponding empirical matrix to calculate and assign branch lengths. This function also randomly resolves polytomies. Matrices were simulated using sim.morpho() in the ‘DispRity’ package (Guillerme 2018). All simulations had a 9:1 binary to three-state character distribution in the matrix, regardless of matrix dimensions. Matrices were simulated under an equal rates Markov model (Lewis 2001) where rates among characters varied based on a predefined lognormal distribution, and each transition within a character occurred at the same rate with equal probabilities. A log-normal distribution was chosen as more appropriate than a gamma distribution as morphological characters are likely affected by hierarchical integration of characters (Wagner 2012). The standard deviation of the log-transformed rate (sd log) was set to 0.5 in all simulations. The mean of the log-transformed rate (mean log) was varied depending on the analysis (described below), as well as the number of simulations generated per run (Table 2). All metrics calculated for the simulated matrices assumed no character state ordering. RHI for the simulations was performed for 100 iterations per run (n = 100).

*Analysis 1: inflating transition rate—*I tested the sensitivity of RHI, CI and RI at detecting homoplasy by using the transition rate of character state change as a proxy for homoplasy in simulated matrices. Higher transition rates lead to increased homoplasy because a greater number of character state transitions may result in independent acquisitions or reversals to the same state (Simmons et al. 2004; Harrison and Larsson 2015). However, the efficacy of increasing transition rates as a proxy for homoplasy will decrease with increased numbers of character states present in a matrix. Moreover, the assumption of homoplasy increasing in tandem with transition rates may not be valid if transitions between states do not occur with equal probability. Therefore, the simulations assume a conservative 9:1 ratio of binary to three-state characters and equal probabilities of character state change directionality. We varied transition rates by employing a range of values for the mean log (*µ*) transition rate and simulating 20 matrices per rate category (Table 2). The range of values (-8 to 2) for *µ* was selected based on trialling multiple scenarios with different ranges of values; this range of values was appropriate for the empirical datasets investigated here, as it covers very low transition rates up to very high rates. The median, 5% and 95% quantiles of RHI, 1 – CI and 1 – RI were plotted against *µ* for four topologies (Table 3.2) which were chosen based on their contrasting matrix dimensions. We also normalised median treelength between 0 and 1 with the following formula:

(7)

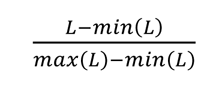

**Table 3.**
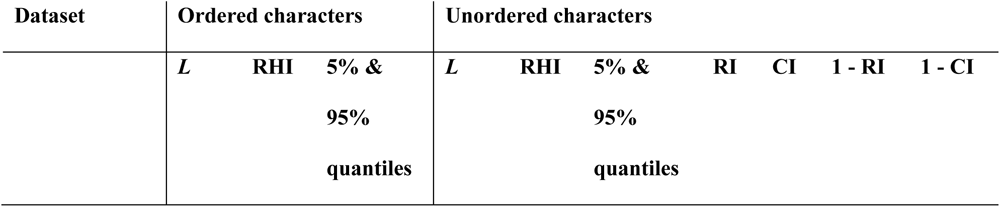
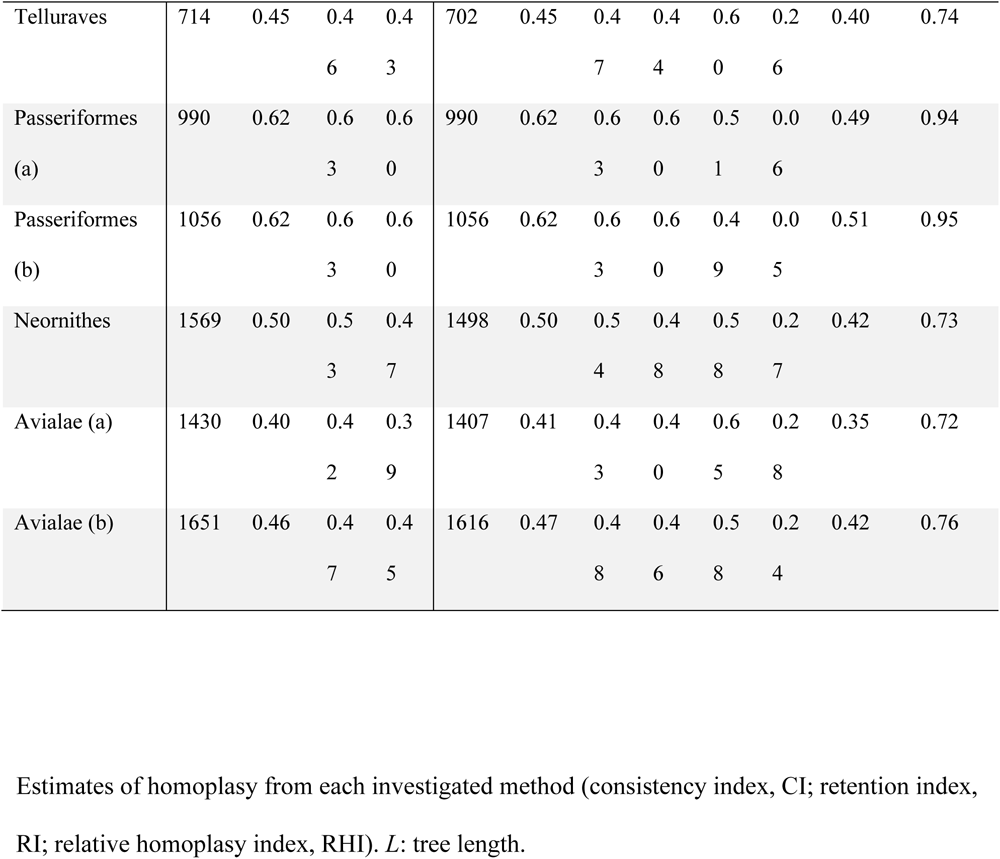
Summary of homoplasy metrics estimated for empirical datasets.

where min(*L*) and max(*L*) represent the minimum and maximum treelengths from the simulation.

*Analysis 2: varying number of taxa with sub-tree sampling—*To test how homoplasy metrics behave with different numbers of taxa, we sampled sub-trees from two empirical topologies (Telluraves and Passeriformes *a*). A new function, subset.trees(), was written to sample sub-trees from a given topology (argument ‘tree’) where the number of tips to be retained (‘ntaxa’) and the number of samples to be taken (‘n’) are specified. One hundred sub-trees were sampled without replacement per set of taxa for each topology, with number of taxa decreasing in increments (Table 3.2). we investigated simulated matrices with two transition rate categories, *µ* = -5 (slow rate) and *µ* = -3 (fast rate). One matrix was simulated for each sub-tree. RHI, 1 – CI and 1 – RI were calculated for each simulated matrix and sub-tree and plotted against number of taxa.

*Analysis 3: varying number of characters—*Similar to Analysis 2, we used the topologies from Telluraves and Passeriformes *a* to investigate metrics with decreasing numbers of characters. Number of characters was specified directly when simulating matrices, and 100 matrices were simulated for each increment of character number, per topology and rate category (Table 2). Rate categories were the same as for Analysis 2. For both Analyses 2 and 3, we performed 20-30 runs per topology and rate category (Table 2).

### Homoplasy in Empirical Datasets

*Analysis 4: comparison of homoplasy metrics across datasets—*RHI, CI and RI were calculated for the six empirical datasets. Topologies were left as they were in the original tree files; therefore, some topologies were not fully bifurcating, and some were unrooted (Fig. 1). All metrics other than RHI were calculated with character states unordered, and RHI was calculated with the set of ordered characters as specified in the original nexus file for each dataset. RHI was run for 1,000 iterations (n = 1,000) per dataset, for both unordered and ordered runs. The 5% and 95% quantiles for RHI were reported alongside the median value.

*Analyses 5 & 6: varying numbers of taxa and characters—*As in Analysis 2, number of taxa was varied for Analysis 5 with sub-trees sampled from all six empirical topologies. Metrics were calculated from the empirical matrices with each sub-tree. Number of characters was varied for Analysis 6 by randomly sub-setting the empirical matrix without replacement. All six datasets were investigated in Analyses 5 and 6. RHI calculations for Analyses 5 and 6 assumed no character state ordering. Between 10–20 increments of taxa/characters were analysed per dataset (Table 2).

## Results

### Simulated Matrices

Analysis 1 evaluated the sensitivity of each metric (RI, CI, and RHI) to increasing character transition rates, a proxy for homoplasy. The results indicate that as transition rate increases (by increasing mean log, *µ*), homoplasy increases (Fig. 2), although the extent of homoplasy detected is dependent on the metric. Normalised tree length (scaled *L*) for each dataset shows that at approximately *µ* = -2, tree length reaches its maximum value for each dataset (Fig. 2). The consistency index rapidly indicates more homoplasy with a small increase in transition rate, and subsequently asymptotes at a much lower *µ* value than the other metrics. In all datasets except Neornithes (Fig. 2c), 1 – CI reaches a maximum value of 1, with Neornithes reaching maximum value at 0.9. Retention index is more robust to increases in transition rate, with 1 – RI reaching an asymptote at approximately equivalent *µ* values to the asymptote point on the X axis of the normalised tree length. However, on the Y axis, 1 – RI does not occupy all homoplasy values between 0 and 1, and instead consistently asymptotes at values below 1. The relative homoplasy index follows the same general pattern as 1 – RI but occupies the full range of homoplasy values between 0 and 1 on the Y axis, and asymptotes at the same point on the X axis (equivalent *µ* values) as the normalised tree length in all datasets.

**Figure 2.**
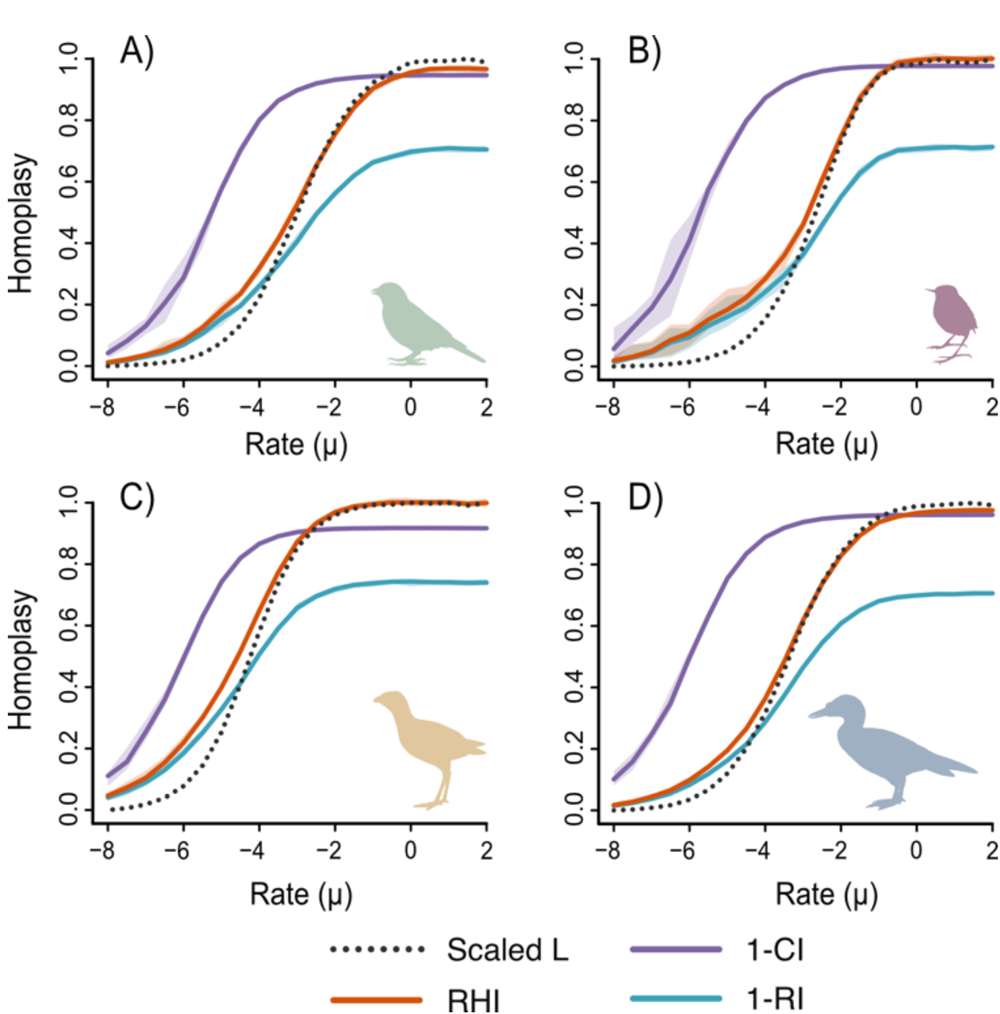
Results from Analysis 1: homoplasy inflation. Simulated matrices using empirical topologies with varying mean log of character state transition rate (µ). Lighter coloured envelopes represent the 5% and 95% quantiles. A) Telluraves topology; B) Passeriformes a topology; C) Neornithes topology; D) Avialae a topology. Scaled L: Tree length normalised between 0 and 1; RHI: relative homoplasy index; 1 – CI: 1 – consistency index; 1 – RI: 1 – retention index.

Analysis 2 tested the sensitivity of metrics to changes in number of taxa, while keeping number of characters the same. Across both low (*µ* = -5) and high (*µ* = -3) transition rates, Analysis 2 indicates that increasing the number of taxa in a topology impacts the homoplasy values estimated by CI, RI and RHI (Fig. 3a, c, e, g). Greater numbers of taxa cause increases in values of 1 – CI, which asymptotes at a maximum value on the Y axis with fewer taxa at higher transition rates (Fig. 3c, g). In contrast, both 1 – RI and RHI values decrease as more taxa are added (Fig. 3a, c, e, g). The extent of decrease in the level of homoplasy detected by 1 – RI and RHI increases markedly at higher transition rates (Fig. 3c, g). In other words, if the character state transition rate is high, a greater decrease in homoplasy will be detected by RI and RHI when taxa are added into a dataset and all other variables stay the same. In Analysis 3, where the number of characters was varied but taxon number was held constant, there is no observable change across the investigated homoplasy metrics as the number of characters was increased (Fig. 3b, d, f, h). More variance was observed for each metric at lower numbers of both taxa and characters, as well as at lower transition rates (Fig. 3).

**Figure 3.**
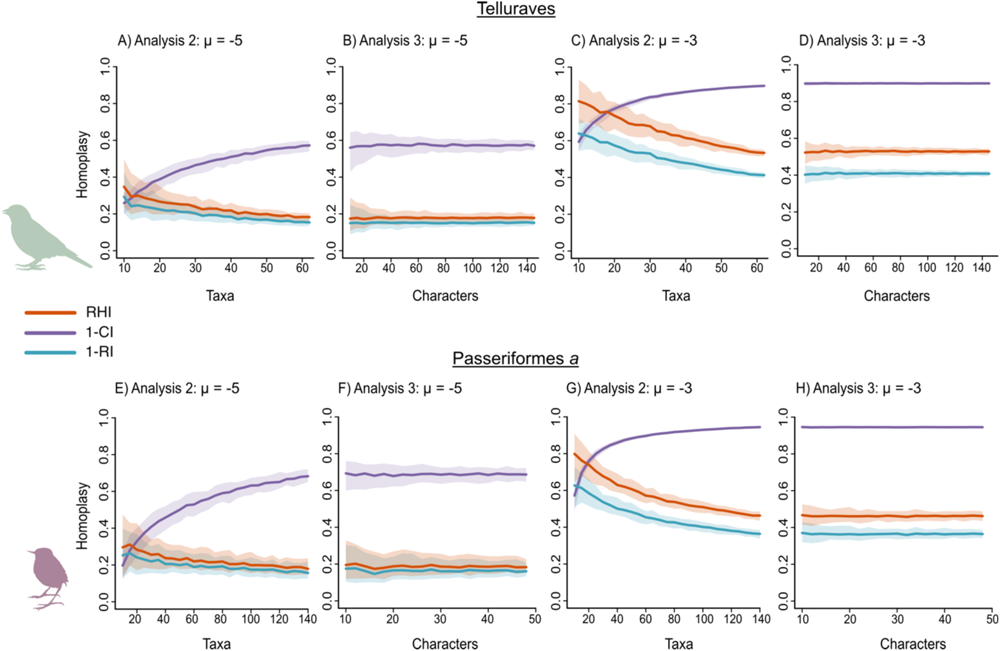
Results from Analyses 2 & 3: varying numbers of taxa and characters in simulated matrices. Simulated matrices using the empirical topologies of Telluraves (A–D) and Passeriformes a (E–F) with varying numbers of taxa (A, C, E, G) and varying numbers of characters (B, D, F, H) at two different mean log values of character state transition rate, slow (µ = -5; A, B, E, F) and fast (µ = -3; C, D, G, H). Envelopes represent the 5% and 95% quantiles of each metric. RHI: relative homoplasy index; 1 – CI: 1 – consistency index; 1 – RI: 1 – retention index.

### Empirical Matrices

In analysis 4, homoplasy was quantified with each metric for each empirical dataset, and results are summarised in Table 3 and Figure 4. Overall, the two Passeriformes datasets show the highest values for all metrics, across both ordered and unordered character treatments, compared to the other bird matrices analysed here. For RHI, the 90% quantile ranges overlap for ordered and unordered treatments for each dataset (Fig. 4). Neornithes shows the greatest 90% quantile range, which is consistent with the comparatively shallow bell curve of the Neornithes *L_null_* distribution (Supplementary Figure 1, Appendix 1).

**Figure 4.**
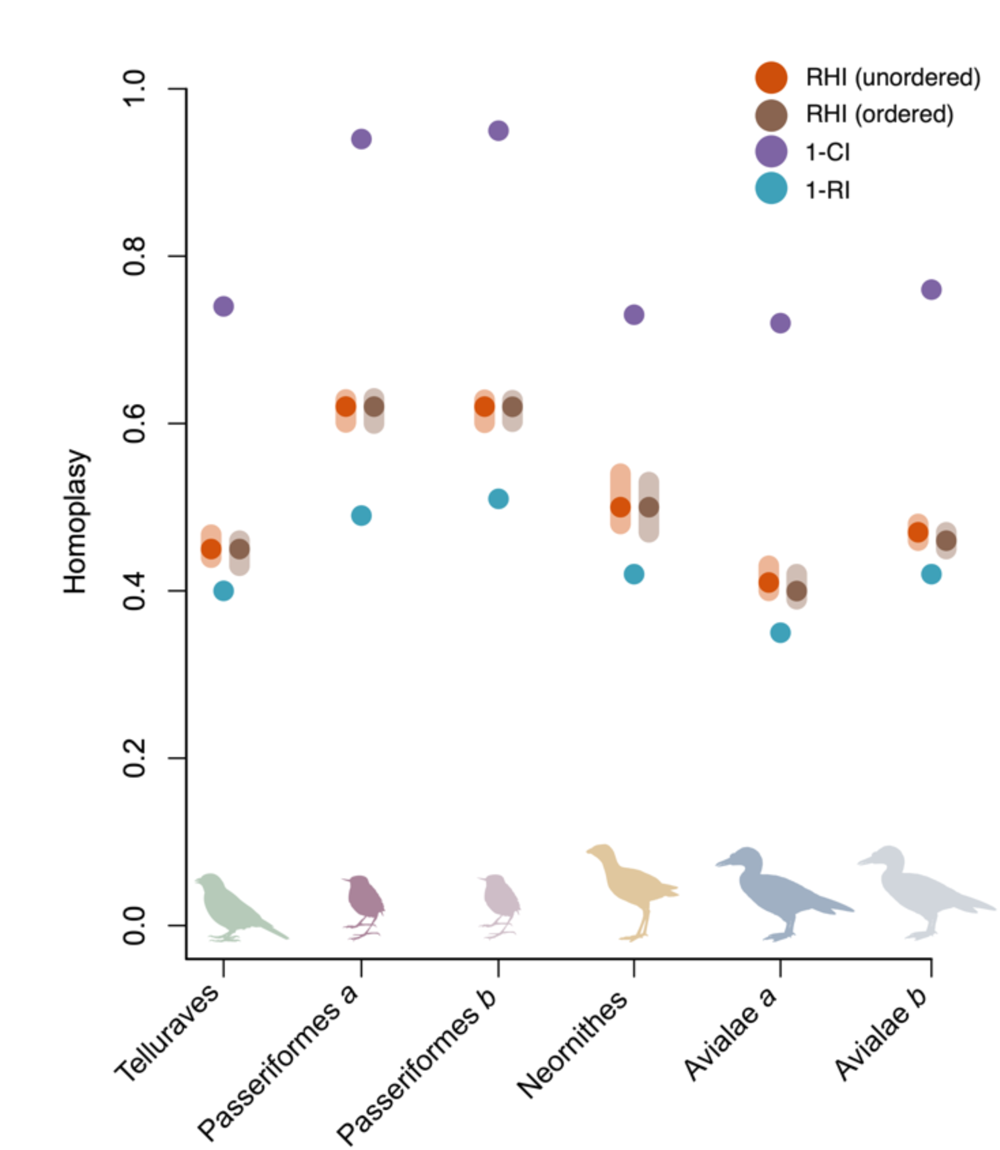
Results from Analysis 4: homoplasy levels in empirical datasets. Summary of all metric results presented in Table 3 for each empirical dataset analysed in this study. RHI envelopes represent 5% and 95% quantile ranges.

Analyses 5 and 6 involved varying the number of taxa and characters, respectively, in each empirical dataset through sampling. Analysis 5 also indicates that with increasing numbers of taxa, 1 – CI increases, but 1 – RI and RHI decrease (Fig. 5a, c, e, g, i, k). However, the manner in which values of 1 – CI, 1 – RI and RHI change with increasing the size of the taxon samples varies depending on the empirical dataset investigated. Overall, 1 – CI changes more abruptly than 1 – RI and RHI with increases in taxon sample, which illustrate comparatively shallow decreases with additional taxa. Similar to Analysis 3, Analysis 6 shows that increasing number of characters does not influence any of the investigated homoplasy metrics (Fig. 5b, d, f, h, j, l).

**Figure 5.**
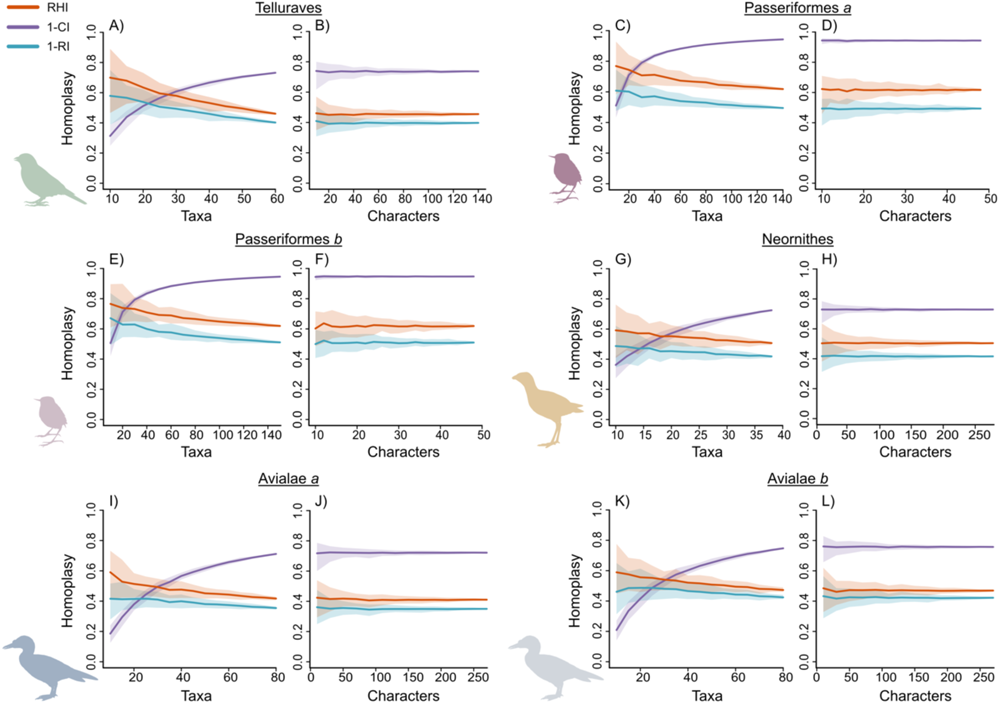
Results from Analyses 5 & 6: varying numbers of taxa and characters in empirical matrices. All empirical datasets with varying numbers of taxa (A, C, E, G, I, K) and varying numbers of characters (B, D, F, H, J, L). Envelopes represent the 5% and 95% quantiles of each metric.

## Discussion

Our new metric, the relative homoplasy index (RHI), uses a novel tip randomisation approach to calculate the relative extent of homoplasy in a phylogenetic character-taxon matrix by comparing the observed amount of homoplasy in that dataset to an estimated null value for that matrix and topology. RHI is computationally tractable as implemented in the *R* statistical environment (github.com/LizzySteell/Homoplasy) and provides better cross-comparable estimates of homoplasy than the commonly used consistency index (CI) and retention index (RI). Through analyses of both simulated and empirical discrete character matrices, we show that RHI is more accurate at detecting homoplasy than CI and more sensitive than RI, particularly in datasets with high levels of homoplasy.

### Patterns of Homoplasy

Although RI has been considered robust with respect to variable numbers of taxa and characters in phylogenetic datasets (Murphy et al. 2021), its major limitations are that it underestimates the proportional extent of homoplasy in a given dataset, and cannot be appropriately compared across alternative datasets because it is not scaled between zero and one, but rather between zero and an arbitrary value conditioned on each particular dataset (Archie 1996). As such, the main advantage of this newly proposed metric, the relative homoplasy index (RHI) over RI is that values of RHI are scaled consistently (Fig. 2), and therefore should provide a more useful method to accurately estimate the relative extent of homoplasy in empirical datasets. The impact of this scaling consistency becomes particularly important when datasets exhibit elevated levels of homoplasy; the simulations in Analysis 1 predict that matrices with greater overall transition rates, and therefore higher levels of homoplasy, will show a greater divergence between 1 – RI and RHI values than matrices with lower transition rates (Fig. 2). In other words, RI underestimates levels of homoplasy to a greater extent in datasets exhibiting greater degrees of homoplasy. As expected, Analysis 4 shows that the empirical matrices with the highest levels of homoplasy (Passeriformes *a* and *b*) show a greater divergence between 1 – RI and RHI than the other datasets (Fig. 4). In addition, RHI values are consistent across the alternative versions of the Passeriformes dataset (Passeriformes *a* and *b*), but the RI values for each differ slightly (Table 3).

Analyses 2 and 5 show that homoplasy as estimated by RHI and RI decrease as taxon numbers increase, whereas homoplasy detected by CI increases considerably with the addition of taxa (Figs. 3 & 5; Table 2). This pattern in CI mirrors early work investigating homoplasy in discrete matrices, which concluded that adding more taxa into a dataset would increase estimated levels of homoplasy (Sanderson and Donoghue 1989). This longstanding hypothesis has been corroborated by numerous subsequent studies (Meier et al. 1991; Hauser and Boyajian 1997; Murphy et al. 2021), but the explanation of why homoplasy increases with the addition of taxa has been somewhat simplistically attributed to the increased detection of individual homoplastic character state changes with the addition of taxa and/or characters (Sanderson and Donoghue 1989; Hauser and Boyajian 1997; Murphy et al. 2021).

CI decreases due to cumulative homoplastic events in a dataset resulting in a greater tree length. However, when considering CI in this format:

(8)

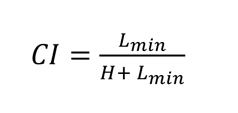

*H* represents the number of extra steps for a tree with a given matrix, and can be considered an unscaled, arbitrary value of homoplasy (Kluge and Farris 1969). *L_min_* does not change with respect to additional taxa, and only varies with changes in number of characters or the composition of character states in a matrix. Therefore, although a tree has a greater probability of gaining homoplastic state changes with the addition of more branches (i.e., increases in *H* via the addition of taxa), this raw quantification of homoplasy in the CI equation is arbitrary and non-comparable; thus, the CI is not a useful tool when attempting to compare estimates of homoplasy from alternative datasets. The original purpose of CI was to measure the consistency of a given dataset with a given tree in order to illustrate how far an empirical topology diverged from a theoretical zero-homoplasy topology derived from a version of the matrix where each character underwent the minimal number of changes (i.e., *L = L_min_*; Kluge and Farris 1969; Farris 1989). The use of CI to quantify homoplasy has been warned against previously due to its high sensitivity towards increasing numbers of taxa (Archie 1989, 1996). We argue that CI should not be considered a measure of homoplasy at all (after all, the opposite of ‘consistency’ as described above does not equate to homoplasy). Rather, CI merely provides a proxy for homoplasy that performs poorly in the face of variable taxon numbers—instead of directly tracking homoplasy, CI tracks an increase in tree length that does not scale proportionally with changes in relative homoplasy.

RI and RHI, on the other hand, are expected to measure a decrease in homoplasy with the addition of taxa to a given dataset, as demonstrated in our results (Figs. 3 & 5). The addition of branches to a topology leads to an increase in the total potential amount of homoplasy in a dataset (*L_max_* in the case of RI and *L_null_* in the case of RHI), due to the exponentially increasing number of possible arrangements of tips as taxa are added (Felsenstein 1978). However, provided the rate distribution of character state transitions for the matrix is maintained, there is no reason to expect more homoplasy in a dataset with more taxa. On the contrary, observed homoplasy values will be proportionally lower compared to the increasing potential homoplasy of the dataset as a taxon sample grows in size.

We find that CI, RI and RHI do not vary with increasing numbers of characters (Figs. 3 & 5). For CI, provided the rate distribution of character state transitions is the same regardless of number of characters, *L* and *L_min_* (and therefore *H*) increase proportionally as characters are added. For RI and RHI, again assuming transition rates are maintained, *L* and *L_max_* and *L* and *L_null_*, respectively, also increase proportionally as characters are added.

Previous studies investigating patterns of detected homoplasy with CI, RI and additional parsimony metrics such as the homoplasy excess ratio (Archie 1989) and the homoplasy slope ratio (Meier et al. 1991) report conflicting results. Archie (1989) observed that RI (therein referred to as the ‘homoplasy excess ratio maximum’) detects less homoplasy in subsets of datasets with more taxa, although a slight increase in homoplasy with the addition of characters is observed (Archie 1989). Other studies using multiple non-subset empirical datasets report mixed results regarding RI. For example, Hauser and Boyajian (1997) found a slight decrease in homoplasy with increasing taxa in their meta-analysis of 600 datasets, but Murphy et al. (2021) showed the opposite trend in an analysis of 364 datasets. These variable patterns are likely explained by idiosyncrasies specific to the datasets analysed; each dataset exhibits a particular amount of estimated homoplasy dependent on factors such as tree shape, character state space (Simmons et al. 2004) and biological drivers of homoplasy. As our results show, larger datasets do not show more homoplasy based on their dimensions alone.

### Future Directions for Homoplasy Quantification

Tree shape is expected to impact patterns of homoplasy estimation (Simmons et al. 2004), which is why a major advantage of the RHI is the tip randomising process. We randomised tips to retain the overall tree shape of each empirical tree, rather than sampling randomly generated trees that may come from a distribution of variable tree shapes with unequal frequencies (Felsenstein 2004). Simmons et al. explored the effect of tree shape on RI and CI with simulated datasets and found that with increasing transition rate, homoplasy detected by RI is affected substantially depending on tree shape. At low transition rates, levels of homoplasy increase significantly more for a completely asymmetrical, unequal branch-length tree than they do for a completely symmetrical, equal branch-length tree (Simmons et al. 2004). Investigating homoplasy patterns with random trees generated by different methods, for example those implemented in ‘APE’ (Paradis et al. 2004), was beyond the scope of the current study but could be useful for the development of new, non-parsimony methods for homoplasy quantification.

In the future, the RHI could be adapted and expanded to estimate homoplasy in a probabilistic framework. Instead of counting steps directly from the matrix, branch-lengths representing evolutionary change that are generated from statistical phylogenetic frameworks (i.e., maximum likelihood or Bayesian inference) could be summed to estimate the observed homoplasy in the dataset. However, randomising the tips would not be a sufficient method to generate a null distribution of summed branch lengths because branch lengths remain the same if the tree shape is retained. Therefore, an alternative way to simulate random trees in order to estimate total potential homoplasy would be required. Statistical trait evolution models, such as the birth-death model (Paradis 2010; Stadler 2011), could be utilised with parameters modelled after the empirical topology to retain its characteristics. This would open new avenues for estimating homoplasy beyond morphological datasets, and could potentially be applicable to molecular sequence data.

The applications of quantifying homoplasy are far reaching. Homoplasy estimates can be used to partition morphological datasets to improve phylogenetic resolution using per-character homoplasy scores to place characters into rate categories (Rosa et al. 2019). It would be possible to use a per-character application for RHI for this, although this step may prove time-consuming. Using RI would likely be sufficient and faster for estimating levels of homoplasy for individual characters in a matrix (for instance, by using the ‘sitewise’ argument in the RI() function in ‘phangorn’ (Schliep 2011)) in order to partition characters into approximate categories. Nonetheless, for accurate estimates of per-character or whole matrix homoplasy, RHI will perform better. Additionally, quantifying homoplasy enables the exploration of why clades are more morphologically labile or constrained. When accompanied by methods investigating character state space and character state exhaustion (Wagner 2000; Hoyal Cuthill 2015; Brocklehurst and Benson 2021), homoplasy quantification can become a tool to better understand patterns in morphological diversification and evolution (Sidlauskas 2008). The RHI metric provides an important starting point in comparative analyses of homoplasy as it presents a non-computationally-demanding method to directly compare the relative extent of homoplasy across empirical phylogenetic datasets.

## Acknowledgements

We thank D. Ksepka for providing a tree file of the published Telluraves topology for analyses. We are indebted to R. Benson for help and support with the methodological approach. We are grateful to D. Brennan, J. Benito, G. Navalón, M.G. Burton, and A. Chen for insightful discussion. This work was supported by an UK Research and Innovation Future Leaders Fellowship MR/S032177/1 to DJF, a Natural Environment Research Council NE/S007164/1 to EMS, and a Swedish Research Council Starting Grant Within Natural and Engineering Sciences ÄR-NT 2020-03515 to AYH. For the purposes of open access, the authors have applied a Creative Commons Attribution (CC BY) license to any Author Accepted Manuscript version arising.

## Supplementary material

Data available from the Dryad Digital Repository: DOI: 10.5061/dryad.34tmpg4rp R code and versions of code arising after this manuscript are available from https://github.com/LizzySteell/Homoplasy

## Appendix 1 Investigating distribution of null tree lengths

**Supplementary Figure 1.**
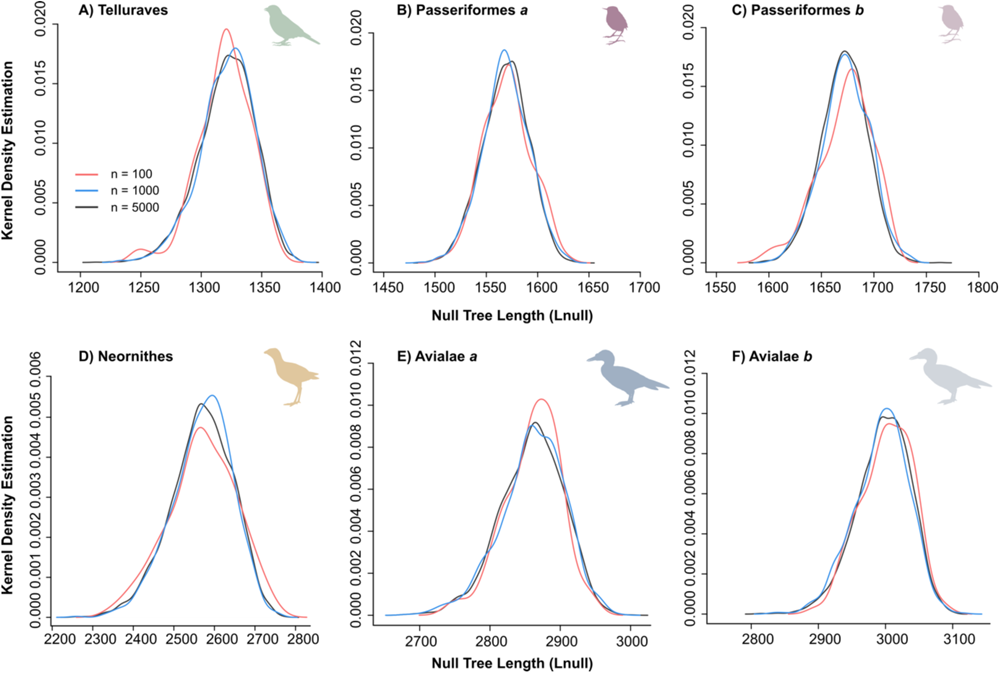
Kernel density plots of null tree length for each empirical dataset. Null tree length distributions during RHI estimation were visualised for different numbers of null trees. Equivalent normal distributions for n = 100, 1,000 and 5,000 suggest 1,000 or even 100 null trees are sufficient to accurately estimate RHI. Kernel density plots were generated using the base R function ‘density()’.

## Literature Cited

1. Archie J.W. 1989. Homoplasy excess ratios: New indices for measuring levels of homoplasy in phylogenetic systematics and a critique of the consistency index. Syst. Zool. 38:253– 269.

2. Archie J.W. 1996. Measures of homoplasy. In: Sanderson M.J., Hufford L., editors. Homoplasy: the recurrence of similarity in evolution. Elsevier. p. 153–188.

3. Benito J., Kuo P.C., Widrig K.E., Jagt J.W.M., Field D.J. 2022. Cretaceous ornithurine supports a neognathous crown bird ancestor. Nature. 612:100–105.

4. Brocklehurst N., Benson R.J. 2021. Multiple paths to morphological diversification during the origin of amniotes. Nat. Ecol. Evol. 5:1243–1249.

5. Cuthill J.F.H., Braddy S.J., Donoghue P.C.J. 2010. A formula for maximum possible steps in multistate characters: Isolating matrix parameter effects on measures of evolutionary convergence. Cladistics. 26:98–102.

6. Farris J.S. 1970. Methods for computing Wagner trees. Syst. Biol. 19:83–92.

7. Farris J.S. 1989. the Retention Index and the Rescaled Consistency Index. Cladistics. 5:417– 419.

8. Felsenstein J. 1978. The number of evolutionary trees. Syst. Zool. 27:27–33. Felsenstein J. 2004. Inferring phylogenies. Sinauer Associates.

9. Field D.J., Benito J., Chen A., Jagt J.W.M., Ksepka D.T. 2020. Late Cretaceous neornithine from Europe illuminates the origins of crown birds. Nature. 579:397–401.

10. Goloboff P.A., Farris J.S., Nixon K.C. 2008. TNT, a free program for phylogenetic analysis. Cladistics. 24:774–786.

11. Guillerme T. 2018. dispRity: A modular R package for measuring disparity. Methods Ecol. Evol. 9:1755–1763.

12. Harrison L.B., Larsson H.C.E. 2015. Among-character rate variation distributions in phylogenetic analysis of discrete morphological characters. Syst. Biol. 64:307–324.

13. Harvey M.G., Bravo G.A., Claramunt S., Cuervo A.M., Derryberry G.E., Battilana J., Seeholzer G.F., Shearer McKay J., O’Meara B.C., Faircloth B.C., Edwards S. V., Pérez-Emán J., Moyle R.G., Sheldon F.H., Aleixo A., Smith B.T., Chesser R.T., Silveira L.F., Cracraft J., Brumfield R.T., Derryberry E.P. 2020. The evolution of a tropical biodiversity hotspot. Science (80-.). 370:1343–1348.

14. Hauser D.L., Boyajian G. 1997. Proportional change and patterns of homoplasy: Sanderson and Donoghue revisited. Cladistics. 13:97–100.

15. Hoyal Cuthill J. 2015. The size of the character state space affects the occurrence and detection of homoplasy: Modelling the probability of incompatibility for unordered phylogenetic characters. J. Theor. Biol. 366:24–32.

16. Kluge A.G., Farris J.S. 1969. Quantitative phyletics and the evolution of anurans. Syst. Biol. 18:1–32.

17. Ksepka D.T., Grande L., Mayr G. 2019. Oldest Finch-Beaked Birds Reveal Parallel Ecological Radiations in the Earliest Evolution of Passerines. Curr. Biol. 29:657–663.e1.

18. Lankester E.R. 1870. II.—On the use of the term homology in modern zoology, and the distinction between homogenetic and homoplastic agreements. Ann. Mag. Nat. Hist. 6:34–43.

19. Lewis P.O. 2001. A likelihood approach to estimating phylogeny from discrete morphological character data. Syst. Biol. 50:913–925.

20. Meier R., Kores P., Darwin S. 1991. Homoplasy slope ratio: a better measurement of observed homoplasy in cladistic analyses. Syst. Zool. 40:74–88.

21. Murphy J.L., Puttick M.N., O’Reilly J.E., Pisani D., Donoghue P.C.J. 2021. Empirical distributions of homoplasy in morphological data. Palaeontology.:pala.12535.

22. Oliveros C.H., Field D.J., Ksepka D.T., Keith Barker F., Aleixo A., Andersen M.J., Alström P., Benz B.W., Braun E.L., Braun M.J., Bravo G.A., Brumfield R.T., Terry Chesser R., Claramunt S., Cracraft J., Cuervo A.M., Derryberry E.P., Glenn T.C., Harvey M.G., Hosner P.A., Joseph L., Kimball R.T., Mack A.L., Miskelly C.M., Townsend Peterson A., Robbins M.B., Sheldon F.H., Silveira L.F., Smith B.T., White N.D., Moyle R.G., Faircloth B.C. 2019. Earth history and the passerine superradiation. Proc. Natl. Acad. Sci. U. S. A. 116:7916–7925.

23. Paradis E. 2010. Time-dependent speciation and extinction from phylogenies: a least squares approach. Evolution (N. Y). 65:661–672.

24. Paradis E., Claude J., Strimmer K. 2004. APE: Analyses of phylogenetics and evolution in R language. Bioinformatics. 20:289–290.

25. Prum R.O., Berv J.S., Dornburg A., Field D.J., Townsend J.P., Lemmon E.M., Lemmon A.R. 2015. A comprehensive phylogeny of birds (Aves) using targeted next-generation DNA sequencing. Nature. 526:569–573.

26. R Core Team. 2021. R: A language and environment for statistical computing..

27. Rosa B.B., Melo G.A.R., Barbeitos M.S. 2019. Homoplasy-based partitioning outperforms alternatives in Bayesian analysis of discrete morphological data. Syst. Biol. 68:657–671.

28. Sanderson M.J., Donoghue M.J. 1989. Patterns of variation in levels of homoplasy. Evolution (N. Y). 43:1781–1795.

29. Schliep K.P. 2011. phangorn: Phylogenetic analysis in R. Bioinformatics. 27:592–593.

30. Sidlauskas B. 2008. Continuous and arrested morphological diversification in sister clades of characiform fishes: A phylomorphospace approach. Evolution (N. Y). 62:3135–3156.

31. Simmons M.P., Reeves A., Davis J.I. 2004. Character-state space versus rate of evolution in phylogenetic inference. Cladistics. 20:191–204.

32. Sookias R.B. 2020. Exploring the effects of character construction and choice, outgroups and analytical method on phylogenetic inference from discrete characters in extant crocodilians. Zool. J. Linn. Soc. 189.

33. Stadler T. 2011. Simulating trees with a fixed number of extant species. Syst. Biol. 60:676– 684.

34. Steell E.M., Nguyen J.M.T., Benson R.B.J., Field D.J. 2023. Comparative anatomy of the passerine carpometacarpus helps illuminate the early fossil record of crown Passeriformes. J. Anat. 242:495–509.

35. Swofford D.L., Maddison W.P. 1987. Reconstructing ancestral character states under Wagner parsimony. Math. Biosci. 87:199–229.

36. Wagner P.J. 2000. Exhaustion of morphologic character states among fossil taxa. Evolution (N. Y). 54:365–386.

37. Wagner P.J. 2012. Modelling rate distributions using character compatibility: implications for morphological evolution among fossil invertebrates. :143–146.

38. Watanabe A. 2016. The impact of poor sampling of polymorphism on cladistic analysis. Cladistics. 32:317–334.

